# High-fat diet-induced obesity differentially alters circadian gene expression across peripheral tissues

**DOI:** 10.64898/2026.05.08.721864

**Authors:** Shota Kawano, Ryusuke Kobayashi, Yuri Watanabe, Ryota Ueno, Takuma Fujimoto, Atsushi Sawada, Daisuke Sawamura, Mitsunori Miyazaki

## Abstract

Circadian rhythms regulate diverse physiological processes, including metabolism, and their disruption has been implicated in metabolic disorders such as obesity. However, the tissue-specific effects of obesity on peripheral circadian clocks remain incompletely understood. Here, we investigated the impact of high-fat diet (HFD)–induced obesity on circadian gene expression in skeletal muscle, liver, and white adipose tissue (WAT). Mice were fed either a regular diet (RD) or HFD for 6 weeks, followed by tissue collection at 4-hour intervals over a 24-hour period. Under RD conditions, key circadian regulators and their downstream targets exhibited robust 24-hour oscillations across all tissues. In contrast, HFD feeding induced distinct, tissue-specific alterations. In the liver, *Per2, Dbp*, and *Rev-erbα* showed phase-advanced expression patterns, whereas in WAT, rhythmic expression was markedly attenuated. Notably, skeletal muscle largely preserved circadian gene expression patterns, indicating relative resistance to HFD-induced circadian disruption. In addition, HFD feeding altered metabolic gene expression in adipose tissue, characterized by reduced *Pgc1α* expression and increased *Leptin* expression. Together, these findings demonstrate that HFD-induced obesity differentially disrupts peripheral circadian clocks in a tissue-specific manner and highlight skeletal muscle as a relatively resilient tissue. These results provide insight into how circadian dysregulation contributes to metabolic abnormalities in obesity.

## Introduction

Circadian rhythms refer to biological cycles with an approximately 24-hour period, originating from the Latin words “circa” (around) and “diem” (day)^1^. In humans, many physiological processes, including sleep-wake cycle^1^, body temperature regulation^2^, hormone secretion^3^, and metabolic processes^4^, are under strict circadian control.

The 24-hour rhythmicity is generated by a molecular mechanism known as the “core clock”, which operates within individual cells^5,6.^ In mammals, circadian timing is regulated by transcriptional feedback mechanisms involving the core factors Brain and Muscle ARNT-Like 1 (BMAL1) and Circadian Locomotor Output Cycles Kaput (CLOCK), which promote the expression of Period (PER) and Cryptochrome (CRY) genes^7,8.^ The PER and CRY proteins subsequently inhibit BMAL1–CLOCK activity, forming a self-regulatory negative feedback loop. BMAL1–CLOCK heterodimers bind to E-box elements in target gene promoters to drive rhythmic transcription^9,10.^ The oscillation of this transcriptional feedback system over approximately 24 hours generates circadian rhythms at the cellular level^11^.

Molecular clock systems are not limited to the central suprachiasmatic nucleus (SCN) but are also independently present in peripheral tissues such as skeletal muscle, liver, and white adipose tissue^12^. These peripheral clocks play essential roles in maintaining metabolic homeostasis and in regulating energy metabolism, and their proper functioning is crucial for human health^13^. Indeed, genetic disruptions in clock genes or disturbances in circadian rhythms have been associated with various pathological conditions, including metabolic, cardiovascular, oncological, and neuropsychiatric disorders^14–16^. For example, mice deficient in Bmal1 or Clock exhibit impaired glucose metabolism, abnormalities in lipid metabolism, and phenotypes resembling metabolic syndrome^17,18.^Furthermore, epidemiological evidence in humans suggests that circadian disruption, such as that caused by shift work or sleep disorders, is associated with an increased risk of malignancies, cardiovascular disease, obesity, and type 2 diabetes^19^. However, the mechanisms linking dysfunction of peripheral clocks to the development of metabolic disorders, including obesity and diabetes, are not yet fully understood.

In this study, we examined how high-fat diet (HFD)-induced obesity affects the circadian expression of core clock genes (*Bmal1, Per2*, and *Cry1*) and their downstream targets, including *D-site-binding protein* (*Dbp*) and *Rev-erbα* (nuclear receptor subfamily 1 group D member 1, Nr1d1), in peripheral tissues such as skeletal muscle, liver, and white adipose tissue.

## Materials and Methods

### Animals

All animal experiments were performed in accordance with institutional guidelines for the care and use of laboratory animals and were approved by the Animal Ethics and Research Committee of the Health Sciences University of Hokkaido (No. 18-041). Male C57BL/6J mice (8 weeks old, purchased from CLEA Japan) were used in this study. Mice were housed under a controlled 12-hour light (07:00–19:00) and 12-hour dark (19:00–07:00) cycle at a constant temperature of 22.0 ± 1.0°C, with *ad libitum* access to food and water. Mice were randomly assigned to two groups: the control group, fed a regular diet (RD; 5% fat content, Oriental Yeast), and the high-fat diet group (HFD; 32% fat content, CLEA Japan). Dietary intervention was continued for six weeks. At the end of the dietary intervention, tissue sampling was conducted at 4-hour intervals over a 24-hour period, covering six time points. The timing of light onset in the animal facility (7:00 AM) was defined as circadian time (CT) 0, and samples were collected every 4 hours (CT0, CT4, CT8, CT12, CT16, and CT20). Three mice (n=3) were sacrificed at each time point for each group, resulting in 18 mice per group (36 mice in total). At each sampling time point, gastrocnemius muscle, liver, and epididymal white adipose tissue (eWAT) were harvested, immediately frozen in liquid nitrogen, and stored at –80°C until further analysis.

### RNA isolation and real-time PCR reaction

Total RNA was extracted from the tissues using TRIzol reagent (Thermo Fisher Scientific, Rockford, IL, USA) according to the manufacturer’s instructions. Residual genomic DNA was removed using TurboDNase (Thermo Fisher Scientific, Rockford, IL, USA). Complementary DNA (cDNA) was synthesized from 0.5 μg of total RNA using PrimeScript RT Master Mix (Takara Bio, Shiga, Japan). Premix Ex Taq (Probe qPCR), SYBR Premix Ex Taq II, and TP850 thermal cycler system (Takara Bio, Shiga, Japan) were used for realtime-PCR. Pre-designed primer and probe sets for *Bmal1* (Mm00500226_m1), *Per2* (Mm00478113_m1), *Cry1* (Mm00514392_m1), *Rev-erbα* (Mm00520708_m1), *Pgc1α* (Mm01208835_m1), *Gapdh* (Mm99999915_g1) genes were purchased from Thermo Fisher Scientific (Rockford, IL, USA). Primers for SYBR assay (5’-3’, forward and reverse, respectively) were designed for *Dbp* (GGAACTGAAGCCTCAACCAATC and CTCCGGCTCCAGTACTTCTCA), *Adiponectin* (TGTTGGAATGACAGGAGCTGAA and CACTGAACGCTGAGCGATACA), and *Leptin* (CAGCCTGCCTTCCCAAAA and CATCCAGGCTCTCTGGCTTCT). The expression levels of each gene were normalized using the 2^−ΔΔCT^ method, referring to the *Gapdh* as an internal control.

### Quantification and statistical analysis

Unpaired two-tailed t-test with Welch’s correction was used for comparisons between two groups. For multiple group comparisons over time, two-way analysis of variance (ANOVA) followed by Šídák’s multiple comparisons test was performed. Data are presented as mean ± standard deviation (SD). Rhythmicity was assessed using JTK_CYCLE implemented in the MetaCycle package in R (version 4.5.1), a non-parametric algorithm for detecting rhythmic components in time-series data. Genes with Benjamini–Hochberg adjusted q-values < 0.05 were considered to exhibit significant rhythmicity. Detailed statistical methods, sample sizes (n), and group comparisons are provided in the corresponding figure legends. All statistical analyses were performed using GraphPad Prism (version 11.0; GraphPad Software, San Diego, CA, USA) and R.

## Results

After 6 weeks of HFD feeding, mice in the HFD group exhibited significant increases in both body weight and eWAT weight compared to the RD group (Figure 1A, B). In addition, during the intraperitoneal glucose tolerance test, a representative HFD-fed mouse showed delayed glucose clearance following glucose loading (Supplemental Figure 1). These results confirm the successful induction of obesity in HFD-fed mice.

**Figure 1.**
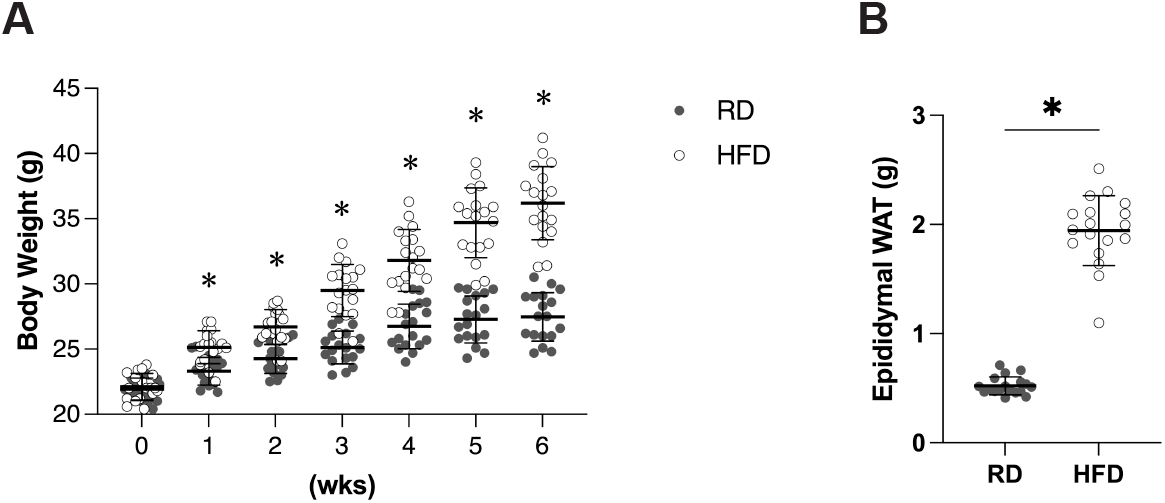
High-fat diet feeding induces obesity in mice. (A)Body weight of mice after 6 weeks of feeding with a regular diet (RD) or high-fat diet (HFD). (B) Epididymal white adipose tissue (eWAT) weight measured at the end of the feeding period. Mice were fed RD or HFD ad libitum for 6 weeks. Data are presented as mean ± SD (n = 18 per group). Statistical significance was determined using an unpaired two-tailed t-test with Welch’s correction. *p < 0.05.

Tissue-specific alterations in circadian gene expression were observed in HFD-fed mice (Figures 2 and 3). Under RD conditions, *Bmal1* and *Per2* exhibited clear 24-hour oscillations with an apparent anti-phase relationship across tissues. JTK_CYCLE analysis confirmed significant rhythmicity of most core clock genes across tissues, although some genes such as Cry1 exhibited weaker rhythmicity in certain tissues (Supplemental Table 1). In the liver, *Per2, Dbp*, and *Rev-erbα* exhibited phase-advanced expression patterns, with a modest reduction in peak expression, particularly for *Dbp* and *Rev-erbα*. In contrast, in eWAT, the oscillatory expression of *Dbp* and *Rev-erbα* was attenuated, with a reduction in amplitude and overall rhythmic robustness, as indicated by JTK_CYCLE analysis. In skeletal muscle, HFD feeding had minimal effects on circadian gene expression, as rhythmic patterns, including phase and amplitude, were largely maintained. These findings indicate that skeletal muscle is relatively resistant to HFD-induced circadian disruption, whereas liver and adipose tissue exhibit more pronounced alterations.

**Figure 2.**
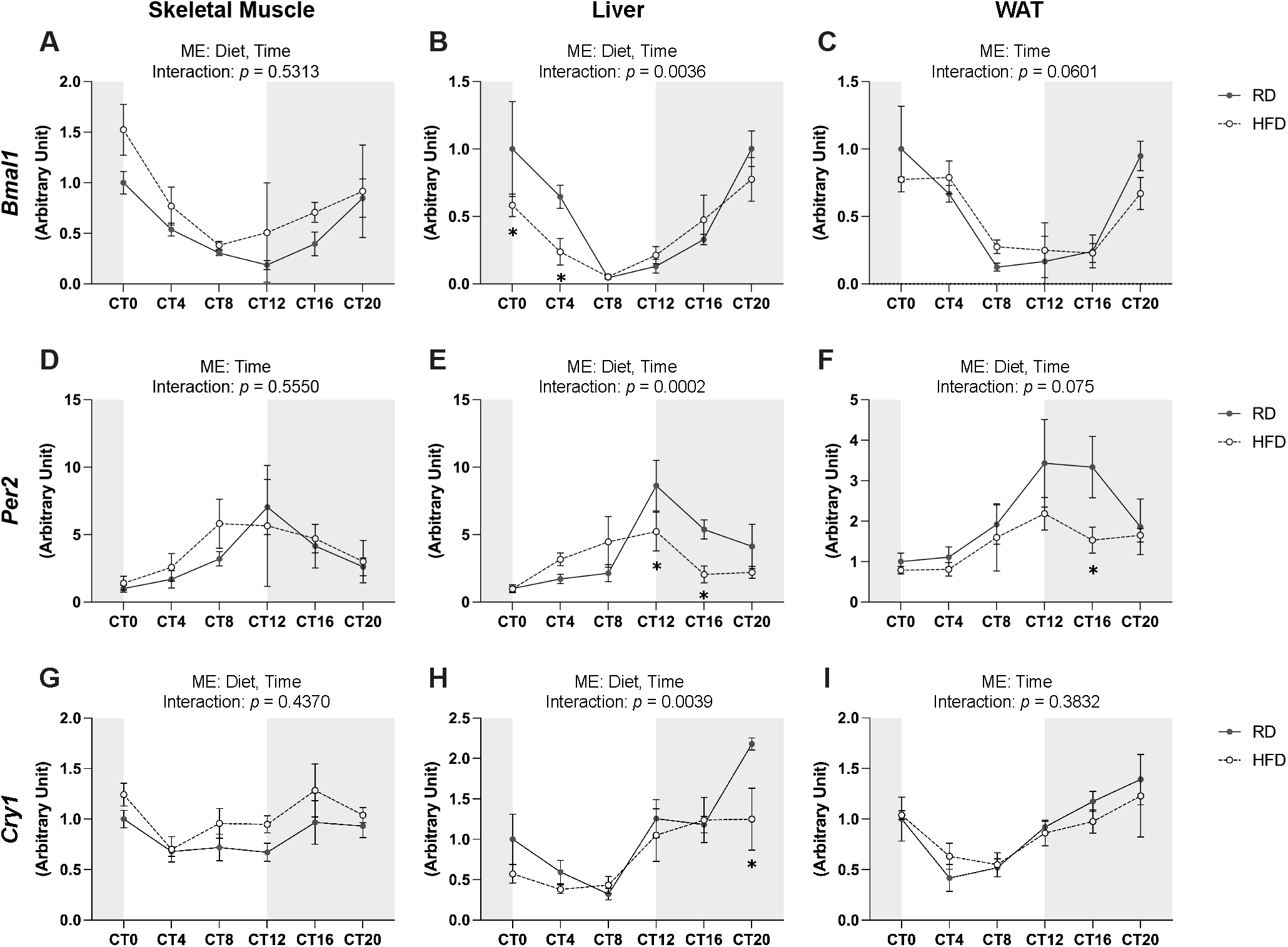
Effects of high-fat diet (HFD) feeding on circadian expression of core clock genes in peripheral tissues. Circadian expression profiles of *Bmal1* (A–C), *Per2* (D–F), and *Cry1* (G–I) in skeletal muscle (A, D, G), liver (B, E, H), and epididymal white adipose tissue (eWAT) (C, F, I) following 6 weeks of feeding with a regular diet (RD) or high-fat diet (HFD). Tissues were collected at 4-hour intervals over a 24-hour period (CT0–CT20). Gray and white backgrounds indicate dark and light phases, respectively. Data are presented as mean ± SD (n = 3 per time point per group). Statistical analysis was performed using two-way ANOVA followed by Šídák’s multiple comparisons test. Main effects (ME) and interaction p-values are indicated in each panel. *p < 0.05 vs. RD at the corresponding time point.

**Figure 3.**
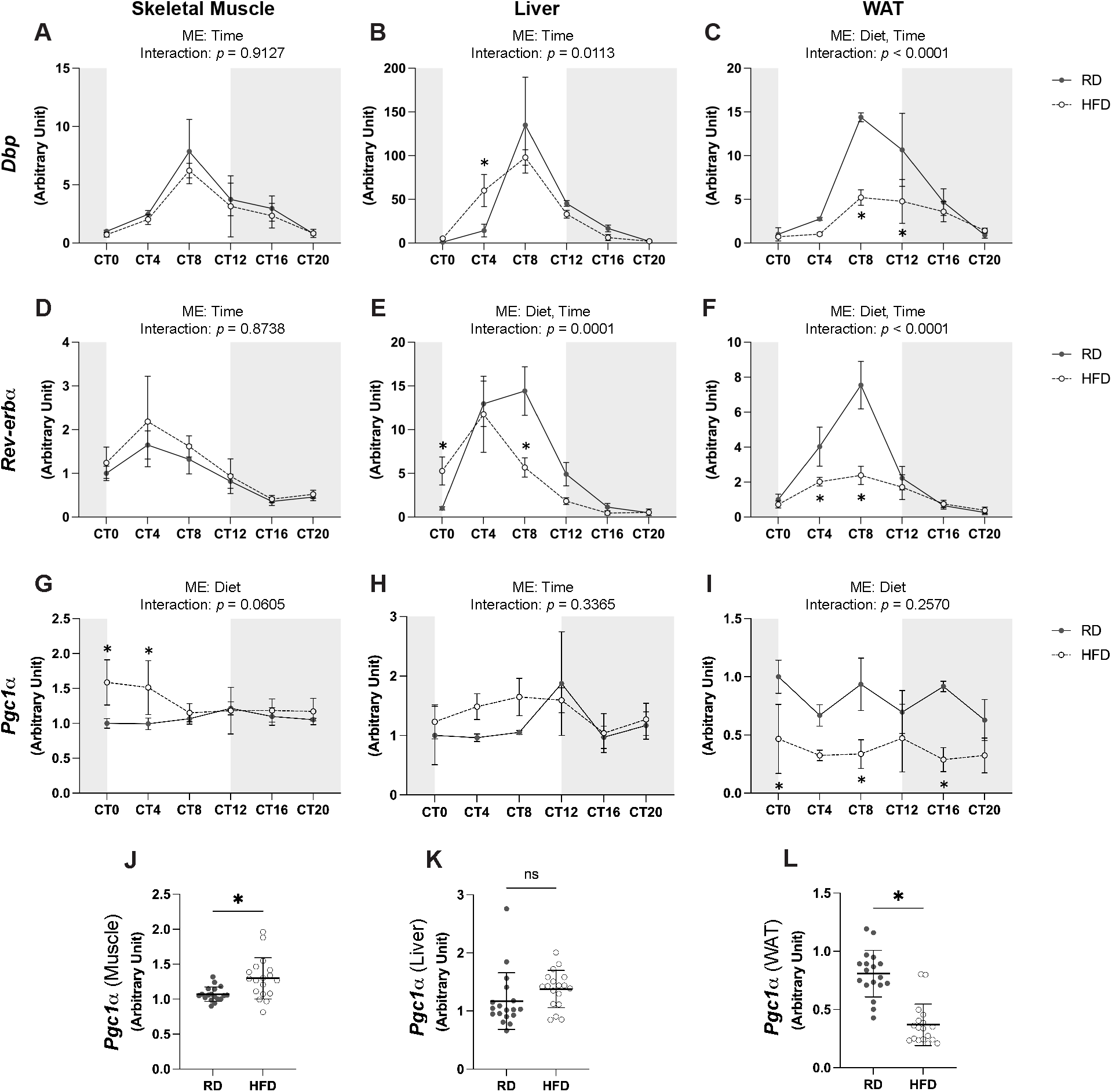
Effects of high-fat diet feeding on circadian expression of clock-controlled and metabolic genes in peripheral tissues. Circadian expression profiles of *Dbp* (A–C), *Rev-erbα* (D–F), and *Pgc1α* (G–I) in skeletal muscle (A, D, G), liver (B, E, H), and epididymal white adipose tissue (eWAT) (C, F, I) following 6 weeks of feeding with a regular diet (RD) or high-fat diet (HFD). Tissues were collected at 4-hour intervals over a 24-hour period (CT0–CT20). Gray and white backgrounds indicate dark and light phases, respectively. Data are presented as mean ± SD (n = 3 per time point per group). Summary comparisons of *Pgc1α* expression between RD and HFD groups are shown in skeletal muscle (J), liver (K), and eWAT (L). Statistical analysis was performed using two-way ANOVA followed by Šídák’s multiple comparisons test for time-course data, and unpaired two-tailed t-test with Welch’s correction for summary comparisons. Main effects (ME) and interaction p-values are indicated in each panel. *p < 0.05 vs. RD at the corresponding time point.

Next, we examined the expression patterns of metabolic genes, including *Pgc1α*, which has been reported to function as a clock-controlled gene and a key regulator of mitochondrial metabolism^20–22^ (Figure 3). In the RD group, *Pgc1α* expression did not exhibit significant circadian rhythmicity in most tissues, although a modest oscillatory pattern was observed in the liver (Figure 3G–I). In contrast, HFD feeding resulted in tissue-specific changes in *Pgc1α* expression, characterized by a significant increase in skeletal muscle, no significant change in the liver, and a marked reduction in eWAT (Figure 3J–L).

Given the marked reduction in *Pgc1α* expression observed in eWAT, we next focused on lipid metabolism–related adipokines (Figure 4). In the RD group, neither *Adiponectin* nor *Leptin* exhibited clear 24-hour rhythmic expression patterns. Under HFD conditions, *Adiponectin* expression showed no consistent changes, whereas *Leptin* expression was markedly elevated across the circadian cycle, indicating increased basal expression levels.

**Figure 4.**
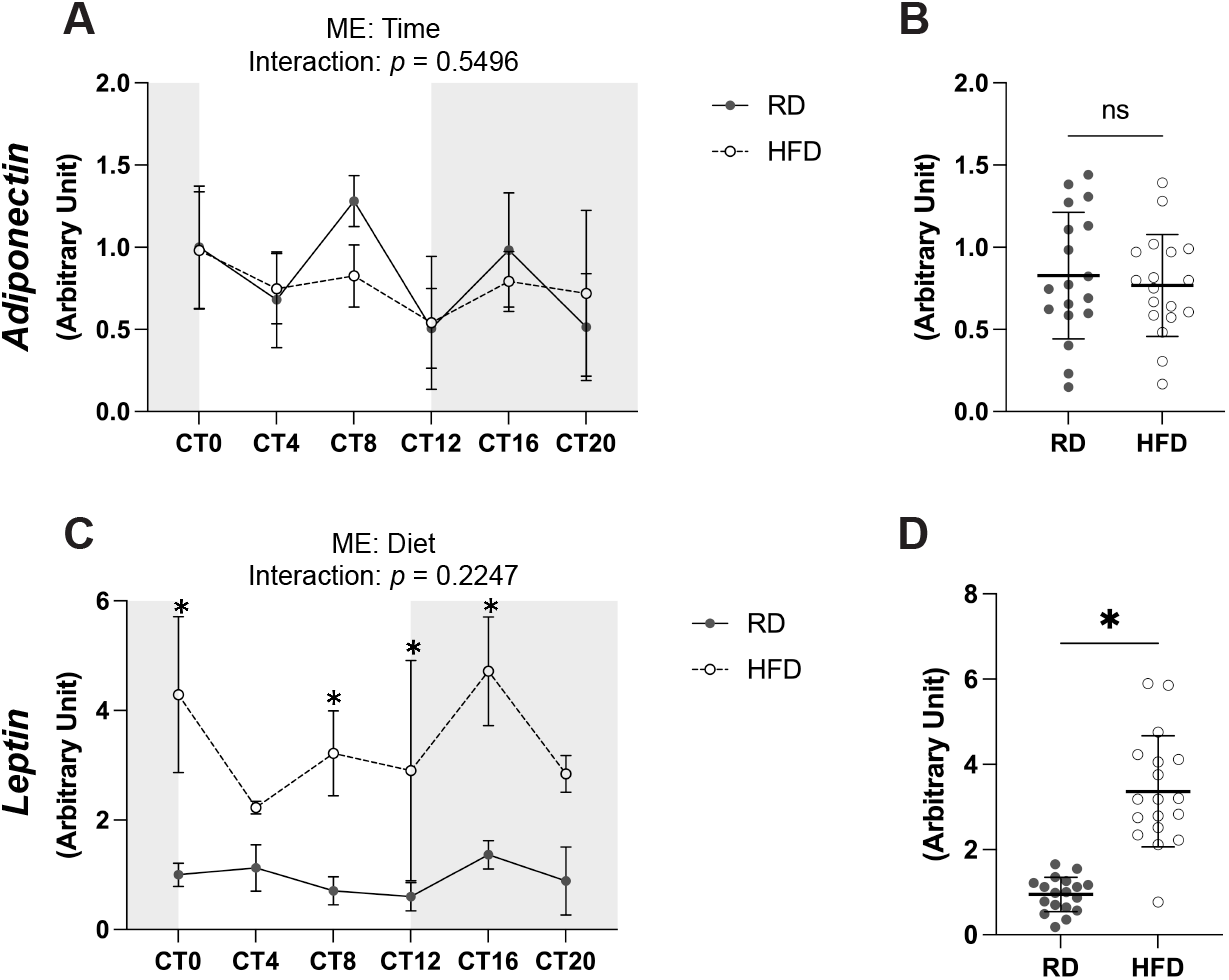
Effects of high-fat diet feeding on adipokine expression in epididymal white adipose tissue. Circadian expression profiles of adiponectin (A) and leptin (C) in eWAT following 6 weeks of feeding with a regular diet (RD) or high-fat diet (HFD). Tissues were collected at 4-hour intervals over a 24-hour period (CT0–CT20). Gray and white backgrounds indicate dark and light phases, respectively. Data are presented as mean ± SD (n = 3 per time point per group). Summary comparisons of adiponectin (B) and leptin (D) expression between RD and HFD groups are shown. Statistical analysis was performed using two-way ANOVA followed by Šídák’s multiple comparisons test for time-course data, and unpaired two-tailed t-test with Welch’s correction for summary comparisons. Main effects (ME) and interaction p-values are indicated in each panel. *p < 0.05.

## Discussion

In this study, we demonstrated that chronic HFD feeding induces tissue-specific alterations in circadian gene expression in peripheral organs. While robust 24-hour oscillations of core clock genes and their downstream targets were observed under RD conditions, HFD feeding led to distinct changes depending on the tissue. Notably, phase-advanced expression patterns were evident in the liver, whereas attenuated rhythmicity, characterized by reduced amplitude, was observed in eWAT. In contrast, skeletal muscle exhibited minimal changes, suggesting that it is relatively less susceptible to HFD-induced circadian disruption.

Accumulating evidence indicates that nutritional factors, including diet composition and feeding patterns, exert critical influences on peripheral circadian clocks. For example, time-restricted feeding can reset peripheral clocks independently of the central SCN, and chronic HFD feeding has been reported to disrupt both behavioral and molecular circadian rhythms^23,24.^ Our findings are consistent with these observations and further extend them by demonstrating that the impact of HFD on circadian regulation is highly tissue-dependent. In the liver, we observed a phase advance in *Per2, Dbp*, and *Rev-erbα* expression, suggesting a shift in the timing of the hepatic clock. The liver is known to be highly responsive to feeding cues and metabolic signals, and thus may be particularly susceptible to HFD-induced perturbations. In contrast, eWAT exhibited a marked attenuation of rhythmic expression, especially in *Dbp* and *Rev-erbα*, indicating a dampening of circadian oscillations. Given the central role of adipose tissue in energy storage and endocrine signaling, such disruption may contribute to metabolic dysregulation under obese conditions.

Interestingly, skeletal muscle largely maintained circadian gene expression patterns, with both rhythmicity and amplitude of core clock genes preserved despite HFD feeding. These findings suggest that skeletal muscle may possess intrinsic mechanisms conferring resilience to metabolic stress–induced circadian disruption. This resilience may be supported by factors such as physical activity–dependent entrainment of muscle clocks^25,26^ and the high metabolic flexibility of skeletal muscle, which enables dynamic adaptation to nutrient availability^21,27.^ In addition, mitochondrial function, closely linked to circadian regulation, has been implicated in maintaining rhythmic stability in peripheral tissues^28^. Although the precise mechanisms remain to be elucidated, our findings highlight a clear tissue-specific hierarchy in susceptibility to HFD-induced circadian alterations.

In addition to alterations in clock gene expression, we observed marked changes in metabolic gene expression, particularly in adipose tissue. *Pgc1α* expression was significantly reduced, consistent with its role in mitochondrial biogenesis and oxidative metabolism^22^. In contrast, *Leptin* expression was constitutively elevated across the circadian cycle, indicating increased basal expression levels, a feature commonly associated with obesity^29^. Given that circadian clocks regulate metabolic gene networks, including the regulation of leptin expression, and that circadian disruption has been linked to leptin resistance and impaired leptin signaling^30,31,^ these findings suggest that dysregulation of peripheral clocks contributes to functional alterations in metabolic pathways. Collectively, our results support a model in which chronic HFD feeding disrupts circadian regulation in a tissue-specific manner, leading to downstream metabolic imbalance, particularly in adipose tissue, which may contribute to the development of obesity-associated metabolic dysfunction^32^.

This study has several limitations. First, the sample size at each time point was limited, which may affect the robustness of rhythm detection. Second, only mRNA expression levels were evaluated; protein expression and functional metabolic outcomes were not assessed. Future studies incorporating protein-level analyses and physiological measurements, such as energy expenditure and locomotor activity, will be necessary to fully elucidate the impact of circadian disruption on metabolic homeostasis.

In conclusion, our results provide evidence that HFD-induced obesity differentially affects peripheral circadian systems and underscore the importance of tissue-specific regulation of circadian rhythms. These findings highlight peripheral clock regulation as a potential target for therapeutic strategies against metabolic disorders.

## Conclusion

In conclusion, our study demonstrates that chronic high-fat diet feeding induces tissue-specific alterations in peripheral circadian gene expression. While the liver exhibits phase-advanced expression patterns and white adipose tissue shows attenuated rhythmicity, skeletal muscle maintains relatively preserved circadian oscillations. In addition, these circadian disruptions are accompanied by changes in metabolic gene expression, particularly in adipose tissue. These findings highlight the importance of tissue-specific regulation of peripheral clocks and provide insight into how circadian dysregulation may be linked to metabolic disturbances in obesity.

## Funding Statement

This work was supported by MEXT KAKENHI Grant Numbers 26K02734, 26H01011, 25K22761, 23K24697, 24H02013, and 24800056 to MM, and by The Uehara Memorial Foundation, The Nakatomi Foundation, and The Takeda Science Foundation. All funding agencies approved the broad elements of study design during the process of grant submission and review and played no direct role in the design of the study, collection, analysis, and interpretation of data, and in writing the manuscript.

## Author Contribution Statement

MM conceived and designed the project; SK, RK, YW, RU, TF, AS, and DS acquired, analyzed, and interpreted the data; SK, RK, and MM wrote, revised, and edited the paper. MM directed the research project.

## Figure Legends

**Supplemental Figure 1. Representative intraperitoneal glucose tolerance test profile**. A representative glucose tolerance curve from an HFD-fed mouse is shown. Mice were fed either a regular diet (RD) or a high-fat diet (HFD) ad libitum for 6 weeks prior to analysis. Mice were fasted overnight, followed by intraperitoneal injection of glucose (2 g/kg body weight). Blood glucose levels were measured at 0, 30, 60, 90, and 120 minutes using a handheld glucometer (Glucocard G, GT-1820; Arkray, Kyoto, Japan). Blood samples were collected from the tail vein via a small incision.

## Notes

### Competing Interest Statement

The authors have declared no competing interest.

