## Supplemental_Figure1 for "High-fat diet-induced obesity differentially alters circadian gene expression across peripheral tissues"

### Supplemental Figure 1

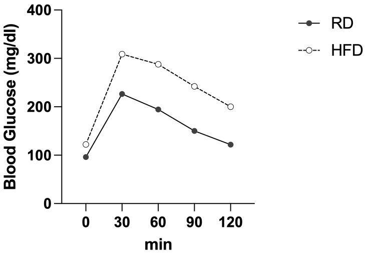


**Supplemental Figure 1. Representative intraperitoneal glucose tolerance test profile.**
A representative glucose tolerance curve from an HFD-fed mouse is shown. Mice were fed either a regular diet (RD) or a high-fat diet (HFD) ad libitum for 6 weeks prior to analysis. Mice were fasted overnight, followed by intraperitoneal injection of glucose (2 g/kg body weight, 20% solution). Blood glucose levels were measured at 0, 30, 60, 90, and 120 minutes using a handheld glucometer (Glucocard G, GT-1820; Arkray, Kyoto, Japan). Blood samples were collected from the tail vein via a small incision.
